# Genetic control of the HDL proteome

**DOI:** 10.1101/405811

**Authors:** Nathalie Pamir, Calvin Pan, Deanna L. Plubell, Patrick M. Hutchins, Chongren Tang, Jake Wimberger, Angela Irwin, Thomas Q. de Aguiar Vallim, Jay W. Heinecke, Aldons J. Lusis

## Abstract

High-density lipoproteins (HDL) are nanoparticles with >80 associated proteins, phospholipids, cholesterol and cholesteryl esters. A comprehensive genetic analysis of the regulation of proteome of HDL isolated from a panel of 100 diverse inbred strains of mice, Hybrid Mouse Diversity Panel (HMDP), revealed widely varied HDL protein levels across the strains. Some of this variation was explained by local, cis-acting regulation, termed cis-protein quantitative trait loci. Variations in apolipoprotein A-II and apolipoprotein C-3 affected the abundance of multiple HDL proteins indicating a coordinated regulation. We identified modules of co-varying proteins and define a protein-protein interaction network describing the protein composition of the naturally occurring subspecies of HDL in mice. Sterol efflux capacity varied up to 3-fold across the strains and HDL proteins displayed distinct correlation patterns with macrophage and ABCA1 specific cholesterol efflux capacity and cholesterol exchange, suggesting that subspecies of HDL participate in discrete functions. The baseline and stimulated sterol efflux capacity phenotypes associated with distinct QTLs with smaller effect size suggesting a multi genetic regulation. Our results highlight the complexity of HDL particles by revealing high degree of heterogeneity and intercorrelation, some of which is associated with functional variation, supporting the concept that HDL-cholesterol alone is not an accurate measure of HDL’s properties such as protection against CAD.

## Introduction

High density lipoproteins (HDL) are composed of a heterogeneous group of lipid-protein complexes that circulate in the blood. HDL-cholesterol levels exhibit strong inverse correlations with coronary artery disease (CAD) in many populations (Castelli and Anderson, 1986; D. J. Gordon and Rifkind, 1989), and their (Kontush et al., 2013; Yetukuri et al., 2010) abilities to promote reverse cholesterol transport and suppress inflammatory responses are consistent with the concept that they protect against the disease (Feig et al., 2014; Tall and Yvan-Charvet, 2015). However, evidence from Genome-Wide Association Studies (GWAS) has suggested that association with CAD may not be causal (Helgadottir et al., 2016). Certain alleles that raise HDL-cholesterol levels at GWAS loci were not associated with protection against CAD in “Mendelian randomization” studies. Additionally, clinical traits with drugs that raise HDL-cholesterol did not show protection against CAD additional to the statin effect. It has been suggested that the discrepancy may be explained by the heterogeneity of HDL (Navab et al., 2011). Thus, it is possible that certain species of HDL, but not others, provide protection against CAD, and that the genetic or drug perturbations failed to impact those mediating the protection. In particular, certain population studies have found that the efficiency of HDL in mediating cholesterol efflux from cells has been associated with decreased incidence of CAD (Khera et al., 2011; 2017; Rohatgi et al., 2014). In addition to CAD, HDL is likely to mediate a variety of immune and regulatory functions (Barter, 2004; S. M. Gordon and Remaley, 2017; Kypreos, 2017). Since these protective functions are likely regulated by the proteins associated with HDL, understanding the regulation of HDL subpopulation’s proteome heterogeneity is an imperative.

Based on size, density, electrophoretic mobility, and protein content of HDL particles, subspecies of HDL have been identified in a variety of species. These include small lipid-poor as well as larger HDL species that contain a large core of cholesteryl esters (Kontush et al., 2013; Yetukuri et al., 2010). In humans, discrete classes of HDL based on size can be identified. In mice, HDL sizes are more continuous (LeBoeuf and Lusis, 1983) and represent one monodisperse peak. The size and levels of HDL vary in both human and mouse populations (Joshi et al., 2016; Pamir et al., 2016). There are clear functional differences associated with the various size classes of HDL. In particular, the small lipid-poor particles are the best acceptors of cholesterol from cells and thus should be particularly important in mediating reverse cholesterol transport, whereas larger particles, associated with proteins such as paraoxonase 1 (PON1) and APOE, are likely to be important in protecting against inflammation. HDL particles containing proteins such as serum amyloid A (SAA) species tend to lack anti-inflammatory properties (Vaisar et al., 2015).

HDL-C levels have a skewed normal distribution in the general population, and the median levels vary by sex and ethnicity. Linkage based studies from the early 1980’s have tried to identify the genetic factors that influence plasma HDL-C levels, but many findings have not been replicated due to polygenic nature of this trait, with contributions from multiple small-effect gene variants. Meta-analyses and GWAS results do, however, support the association of HDL-C with variation in CETP, LIPC, LPL, ABCA1, endothelial lipase (LIPG) and LCAT (Hegele, 2009; Thompson et al., 2008). Multiple genetic factors could be present in an individual, creating a polygenic network of HDL-C determinants (Cohen et al., 2004). These determinants include monogenic effectors, such rare homozygous mutations in ABCA1, LCAT, and APOA1 causing extremely low HDL-C (Brooks-Wilson et al., 1999; Kuivenhoven et al., 1996; Ng et al., 1994), and rare homozygous mutations in CETP, LIPC, and SCARB1, causing extremely elevated HDL-C. The mouse models of these variants have been supportive of the human findings (Wang and Paigen, 2005) (Wang & Paigen, 2005). Polygenic determinants have been recently investigated using targeted next-generation sequencing in patients with extremely low and high HDL-C. About 30% of individuals at the extremes of HDL-C had rare large effect and common small effect variants explaining the trait (Dron et al., 2017). Whereas the genetic determinants of plasma HDL-C levels have been well studied, the genetic determinants of HDL proteome and lipidome have never been previously investigated.

To better define the various species of HDL at the level of protein composition and to understand their genetic regulation we used the Hybrid Mouse Diversity Panel (HMDP) with a systems biology approach (Bennett et al., 2010; Ghazalpour et al., 2012). HMDP is a collection of 100 classical laboratory and recombinant inbred (RI) strains that have been genotyped at 135,000 SNPs (Bennett et al., 2010). HMDP provides a confined genetic space (Bennett et al., 2010) that rely on naturally occurring genetic variation that perturbs protein abundance. We performed systems genetics approach using analytical approaches that include genome-wide association, expression quantitative trait locus (eQTL) discovery, functional outcomes and network analysis (Bennett et al., 2010).

We identified HDL proteome and HDL function for each strain. Using quantitative trait locus (QTL) analyses, we performed genetic mapping. First, associations between SNPs and HDL protein levels and function were determined. Second, the effects of SNPs on gene regulation were determined by eQTL analysis using hepatic and adipose tissue-specific gene expression. Based on the genetic variation in HDL observed, we were able to identify numerous genetic factors mediating HDL composition and provide an approximation of the nature of the interactions between the proteins. Finally, we used protein-protein interaction cluster analyses to build an HDL model that describes the group of proteins that constitute the core proteins and the ones that are peripheral. We identified moderate QTLs associated with HDLs sterol efflux capacity. Our results reveal a great deal of heterogeneity and inter-correlation, some of which is associated with functional variation, supporting the concept that HDL-cholesterol alone is not an accurate measure of HDL’s protective properties in terms of CAD.

## Results

### Quantitation of HDL associated protein levels in a panel of 100 inbred strains of mice

HDL isolated from 93 strains of HMDP was subjected to LC-MS/MS (Pamir et al., 2016). This list of robustly identified proteins, ranked by average abundance, is presented in Table 1. A full list of the proteins with peptide spectra matches (PSMs) per biological replicate is presented in Supplemental Table 1.

**Table 1.** List of proteins detected in mouse HDL across HMDP.

| **Gene symbol** | **Gene name** |
| --- | --- |
| Abpa7 | secretoglobin, family 1B, member 7 |
| Ahsg | alpha-2-HS-glycoprotein |
| Alb | albumin |
| Antxr1 | anthrax toxin receptor 1 |
| Antxr2 | anthrax toxin receptor 2 |
| Apoa1 | apolipoprotein A-I |
| Apoa2 | apolipoprotein A-II |
| Apoa4 | apolipoprotein A-IV |
| Apoa5 | apolipoprotein A-V |
| Apob | apolipoprotein B |
| Apoc1 | apolipoprotein C-I |
| Apoc2 | apolipoprotein C-II |
| Apoc3 | apolipoprotein C-III |
| Apoc4 | apolipoprotein C-IV |
| Apod | apolipoprotein D |
| Apoe | apolipoprotein E |
| Apoh | apolipoprotein H |
| Apom | apolipoprotein M |
| Apon | apolipoprotein N |
| Arsg | arylsulfatase G |
| Bpifa2 | BPI fold-containing family A member 2 |
| C3 | complement component 3 |
| C4b | complement component 4B (Chido blood group) |
| C4bpa | C4b-binding protein |
| Camp | cathelicidin antimicrobial peptide |
| Cd97 | CD97 antigen |
| Clec14a | C-type lectin domain family 14, member a |
| Clu | clusterin |
| Cst6 | cystatin E/M |
| Ctsd | cathepsin D |
| Dmkn | dermokine |
| Egfr | epidermal growth factor receptor |
| F10 | coagulation factor X |
| Fga | fibrinogen alpha chain |
| Fgb | fibrinogen beta chain |
| Fgg | fibrinogen gamma chain |
| Gc | group specific component |
| Gm5938 | predicted gene 5938 |
| Gm94 | predicted gene 94 |
| Gpld1 | glycosylphosphatidylinositol specific phospholipase D1 |
| Grn | granulin |
| H2-Q4 | histocompatibility 2, Q region locus 4 |
| H2-Q10 | histocompatibility 2, Q region locus 10 |
| Hba | hemoglobin alpha chain complex |
| Hbb-b1 | hemoglobin, beta adult major chain |
| Hbb-b2 | hemoglobin, beta adult minor chain |
| Hbbt1 | beta-globin |
| Icam1 | intercellular adhesion molecule 1 |
| Ifi27l2b | interferon, alpha-inducible protein 27 like 2B |
| Igfals | insulin-like growth factor binding protein, acid labile subunit |
| Ighm | Immunoglobulin heavy constant mu |
| Ihh | Indian hedgehog |
| Itgb1 | integrin beta 1 (fibronectin receptor beta) |
| Lcat | lecithin cholesterol acyltransferase |
| Mug1 | murinoglobulin 1 |
| Napsa | napsin A aspartic peptidase |
| Obp1a | odorant binding protein 1a |
| Pcyox1 | prenylcysteine oxidase 1 |
| Pf4 | platelet factor 4 |
| Plg | plasminogen |
| Pltp | phospholipid transfer protein |
| Pon1 | paraoxonase 1 |
| Pon3 | paraoxonase 3 |
| Ppic | peptidylprolyl isomerase C |
| Psap | prosaposin |
| Rab1a | ras-related protein Rab-1a |
| Rbp4 | retinol binding protein 4, plasma |
| Saa1 | serum amyloid A 1 |
| Saa2 | serum amyloid A 2 |
| Saa4 | serum amyloid A 4 |
| Scgb1b2 | secretoglobin, family 1B, member 2 |
| Sell | selectin, lymphocyte |
| Serpina1a | serine (or cysteine) peptidase inhibitor, clade A, member 1A |
| Serpina1e | serine (or cysteine) peptidase inhibitor, clade A, member 1E |
| Serpina3k | serine (or cysteine) peptidase inhibitor, clade A, member 3K |
| Serpinc1 | serine (or cysteine) peptidase inhibitor, clade C (antithrombin), member 1 |
| Tf | serotransferrin |
| Tfpi | tissue factor pathway inhibitor |
| Tfrc | transferrin receptor |
| Ttr | transthyretin |
| Vcam1 | vascular cell adhesion molecule 1 |
| Vtn | vitronectin |

We used two common ways to normalize the PSMs: 1) normalized each PSM to spiked yeast Carboxypeptidase Y protein (Carvalho et al., 2008; Liu et al., 2004) (Liu *et al*, 2004; Carvalho *et al*, 2008), 2) normalized every PSM to observed total PSMs (Vaisar et al., 2007). We present findings from both analyses: Main text data and figures are from yeast normalized data analysis, whereas the total normalized analysis is presented in the supplements. Proteins that were identified with at least 2 unique peptides were subjected to further analysis. Of these proteins, 34 were shared across all strains (Figure 1 and Supplemental Figure 1). The proteome was analyzed for proteins represented in strains by quintiles. The bottom quintile proteins, identified in less than 20% of the 93 strains, were discarded. The remaining 155 proteins identified were used perform the QTL analyses (Figure 2 and Supplemental Figure 2). To identify how the proteins correlated with each other and with the functional metrics of HDL, we used proteins that were represented in 80% of the strains (81 proteins total).

**Figure 1:**
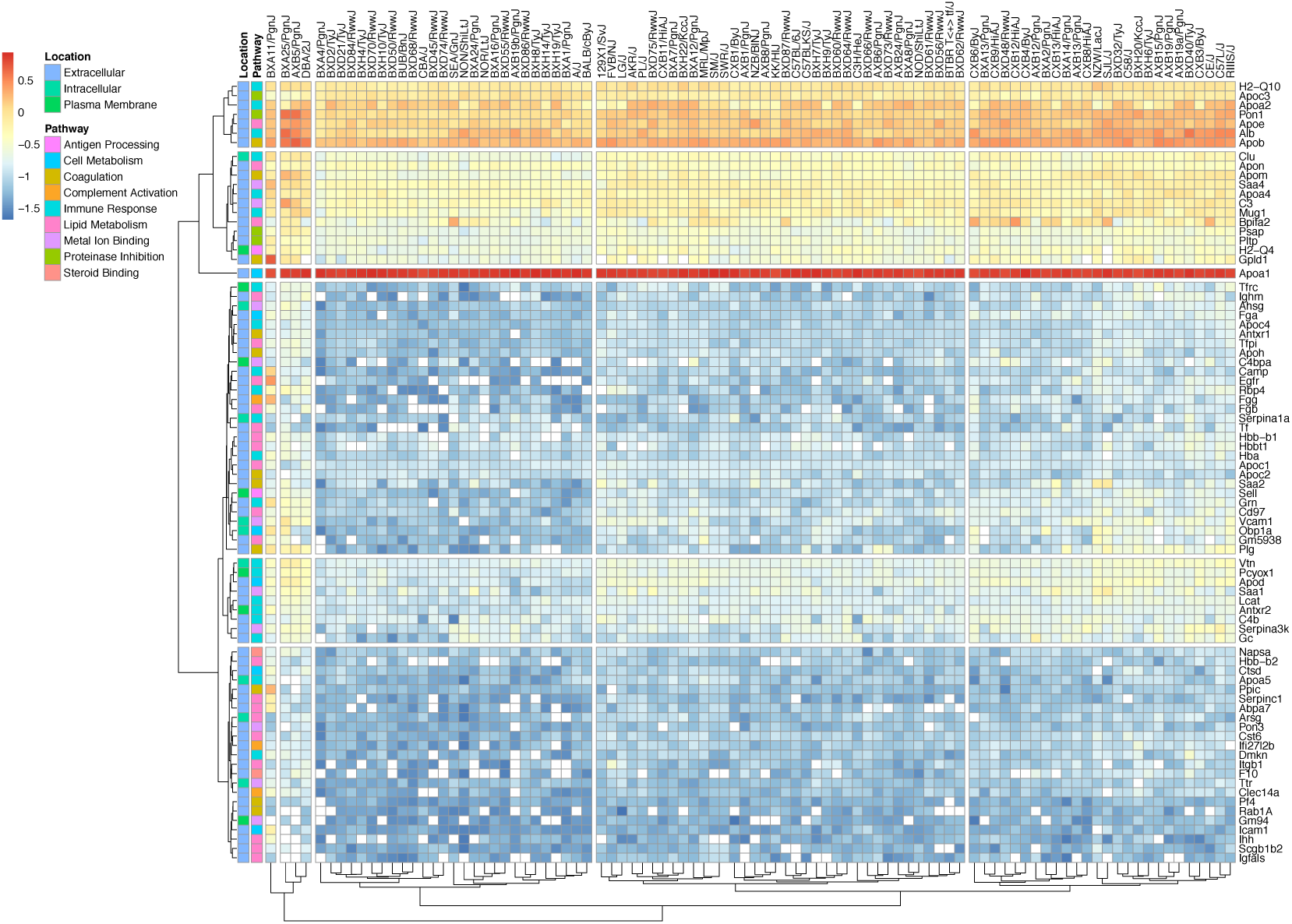
The heatmap visualization of the HDL protein abundances across 93 strains. The proteins, their biological functions and cellular locations are represented. Logarithmic transformation of the yeast normalized data has been performed to accommodate the abundance distribution form high (red) to low (dark blue). White squares represent not available values. Both the proteins and the strains were clustered using Euclidean distances.

**Figure 2.**
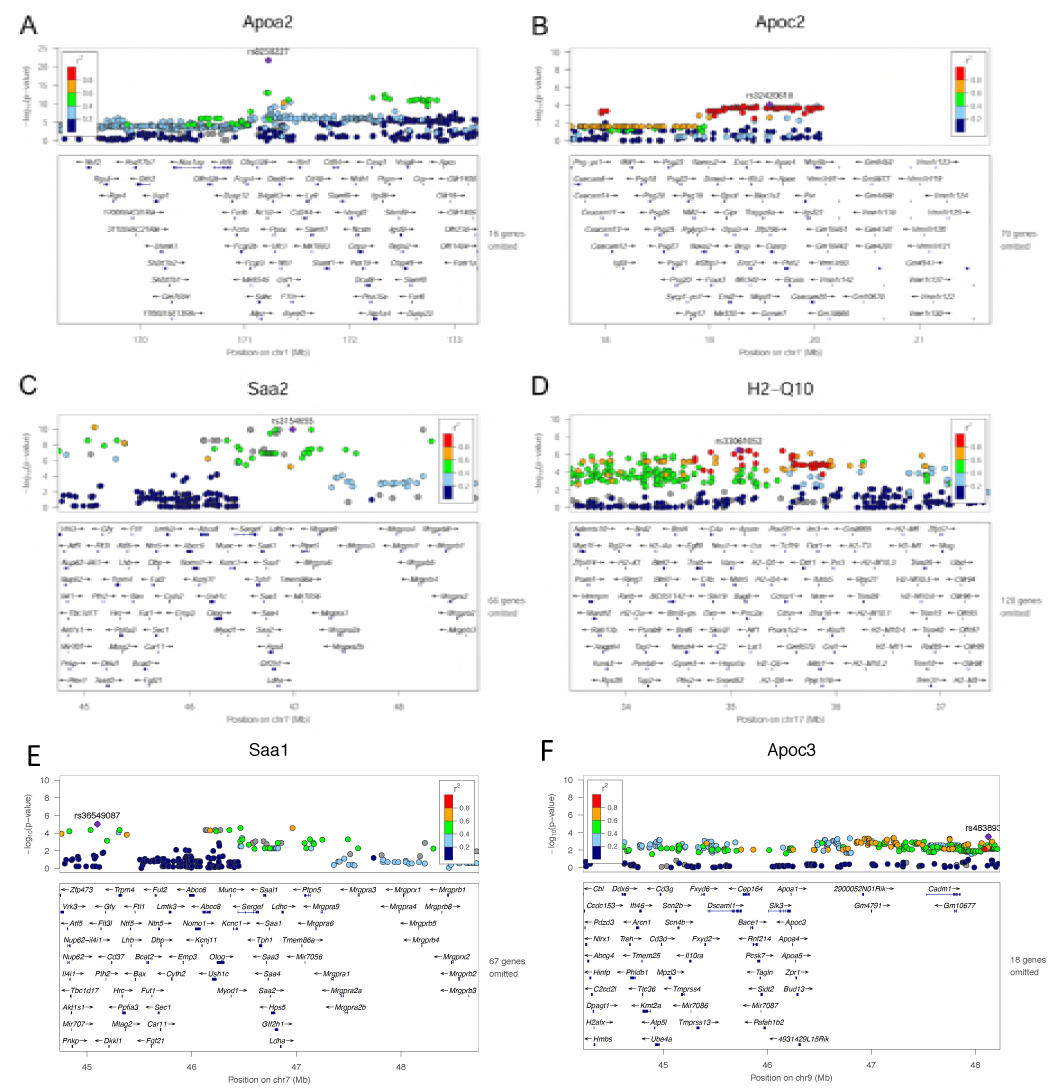
A representative QTLs for HDL proteins. Loci associated with protein levels of APOA2, B), APOC2, C) SAA2, D) H2-Q10 E)SAA1 F)APOC3

The heatmap representation of the relative abundance of proteins across 93 strains, the biological processes in which they participate and their cellular locations are visualized to provide information about the relative abundance and their spatial (Euclidean) clustering (Figure 1 and Supplemental Figure 1). We have curated outputs from publicly available enrichment databases such as DAVID, PANTHER, KEGG, and Gene Ontology to classify the proteins according to the biological functions in which they participate and the cellular location they are most likely to operate. As expected most of the proteins (65/81) were extracellular. Our results are in agreement with previous reports (Vaisar et al., 2007) and show that HDL is associated with proteins that play a role in antigen processing, cell metabolism, coagulation, complement activation, immune response, lipid metabolism, metal ion binding, proteinase inhibition, and steroid binding. All of the proteins identified are replicated by previous studies as shown by HDL proteome watch website (http://homepages.uc.edu/∼davidswm/HDLproteome.html) by Dr. Sean Davidson’s Laboratory at University of Cincinnati.

We had previously shown in a smaller study with five inbred strains that HDL proteome predicts the genealogy of the strains (Pamir et al., 2016) suggesting a hereditary component. The heritability of the HDL proteome was estimated by calculating the broad-sense heritability scores (H2) for each protein (Supplemental Table 1). Broad-sense heritability (H2) captures the phenotypic variation due to genetic factors such as dominance and epistasis (Visscher et al., 2008). The HMDP consists of 29 “classic” inbred strains and about 70 recombinant strains derived from 5 different sets of parental strains (BxA or AxB, BxD, CxB, and BxH) (Bennett et al., 2010). Our cohort has N=1-5 biological replicates per strain. The proteins identified are not always represented in each sample leading to variable N. Despite the incrementally variable genetic canvas of the strains and the technical variability in mass spectrometry analyses 66/155 yeast normalized proteins had a H2 score between 0.10 and 0.90 indicating up to 90% heritability due to genetic factors. The HDL proteome is composed of core set of proteins (∼40) that are detected in every study across the diverse laboratories and sample sets, and a subset of proteins that are acutely regulated by the environment (ie. Diet and inflammation-http://homepages.uc.edu/∼davidswm/HDLproteome.html). The H2 scores for such proteins are expected to be zero. Furthermore, due to our study design, the proteins that are strain specific are also expected to have a H2 of zero.

### Genetic regulation of HDL proteome

Loci contributing to variations in protein levels (protein QTLs or pQTLs) or hepatic transcript levels of the proteins (expression QTLs or eQTLs) were mapped using the FAST LMN, an association algorithm with a mixed model component to correct for population structure. Association analysis was performed using about 200,000 informative SNPs (Bennett et al., 2010), spaced throughout the genome (Table 2). Hepatic transcript levels were from a previous survey of the HMDP maintained on the same chow diet as in this study (Bennett et al., 2010). Loci averaged 500 kb to 2 Mb in size and in most cases contained 1 to 20 genes within a linkage disequilibrium (LD) block, an improvement of more than an order of magnitude as compared to traditional linkage analysis in mice (typically a resolution of 10 to 20 Mb) (Flint et al., 2005). Loci mapping within 1Mb of the gene are termed “local” QTL, suggesting that they probably act in *cis.* For example, promoter or enhancer variants would act in *cis*. Loci mapping greater than 1Mb from the gene of interest are termed “distal”, implying that they act in *trans*, presumably mediated by a diffusible factor such as a transcription factor (Table 2 and Supplemental Table 2). We applied a significance filter of *P*=10^−3^ and 10^−6^ to identify suggestive cis and trans QTLs respectively.

**Table 2.** Local hepatic eQTL and HDL protein pQTL Local eQTL and pQTL with p < 0.001 are listed along with SNP identifiers and their locations

|  |  | Local eQTL |  |  | Local pQTL |  |  |
| --- | --- | --- | --- | --- | --- | --- | --- |
| Gene symbol | gene location | rsID | location | pvalue | rsID | location | pvalue |
| Antxr1 | chr6:87133854-87335775 | rs37023898 | chr6:87529811 | 1.81E-04 | - | - | - |
| Antxr2 | chr5:97884688-98030962 | rs33642547 | chr5:97956955 | 7.35E-09 | - | - | - |
| Apoa2 | chr1:171225054-171226379 | rs46114424 | chr1:170988751 | 1.28E-04 | rs8258227 | chr1:171225758 | 1.63E-22 |
| Apoc2 | chr7:19671584-19681423 | rs32193511 | chr7:19542617 | 2.54E-04 | rs32420618 | chr7:19571593 | 7.92E-05 |
| Apoc3 | chr9:46232933-46235636 | - | - | - | rs48945377 | chr9:46673334 | 5.41E-04 |
| Apoh | chr11:108343354-108414396 | rs29467866 | chr11:107673524 | 8.33E-16 | - | - | - |
| Apom | chr17:35128997-35132050 | rs33061052 | chr17:35078160 | 4.57E-11 | - | - | - |
| Apon | chr10:128254131-128255901 | rs29339020 | chr10:128283501 | 2.27E-11 | - | - | - |
| Arsg | chr11:109473374-109573330 | rs27052095 | chr11:109531899 | 6.33E-07 | - | - | - |
| B2m | chr2:122147686-122153083 | rs27439233 | chr2:121957890 | 1.02E-14 | - | - | - |
| Clu | chr14:65968483-65981548 | rs31035575 | chr14:66419322 | 5.69E-04 | - | - | - |
| Dmkn | chr7:30763756-30781066 | rs8236440 | chr7:30734739 | 5.42E-04 | - | - | - |
| Gpld1 | chr13:24943152-24990753 | rs30001837 | chr13:25375000 | 1.63E-05 | - | - | - |
| H2-Q10 | chr17:35470089-35474563 | rs33079924 | chr17:35760448 | 7.03E-54 | rs33061052 | chr17:35078160 | 2.99E-07 |
| Itgb1 | chr8:128685654-128733200 | rs33614001 | chr8:128711717 | 1.73E-04 | - | - | - |
| Pltp | chr2:164839518-164857711 | rs27317491 | chr2:164014231 | 5.33E-06 | - | - | - |
| Pon1 | chr6:5168090-5193946 | rs30319905 | chr6:4188552 | 9.52E-06 | - | - | - |
| Saa1 | chr7:46740501-46742980 | rs32054038 | chr7:47248355 | 4.11E-04 | rs3154655 | chr7:46985523 | 1.19E-05 |
| Saa2 | chr7:46751833-46754314 | rs32054038 | chr7:47248355 | 1.53E-06 | rs3154655 | chr7:46985523 | 1.09E-10 |

A total of 19 HDL proteins showed significant evidence of local regulation of hepatic transcript levels or protein levels (Table 2). With the exception of *Apoc3*, all of the significant pQTL also exhibited significant eQTL, indicating that genetic variation in protein levels was largely due to regulation of gene expression. In the case of *Apoc3*, while there was no significant eQTL in liver, there was a highly significant eQTL in adipose tissue (*P*=1.8 x 10^−7^) (Supplemental Table 3). *Apoa2* and *Saa2* exhibited much more significant pQTL than eQTL (*P*=1.324e-22, effect size = −0.413 vs. *P*=1.276e-4, effect size = 0.0210 for Apoa2, and *P*=2.559e-10, effect size = 0.0780 and *P*=*1.533e-6, effect size = 0.780 for Saa2 respectively)*, suggesting that the pQTL were due to coding rather than regulatory variations. In the case of *Apoa2*, our previous studies indicated that structural variation affecting translation efficiency was largely responsible for the differences in protein levels among several common inbred strains (Bennett et al., 2010). Also, common coding variants are present among the HMDP strains for both *Apoe3* and *Saa2* (www.informatics.jax.org). As shown in Table 2, many variants significantly affecting HDL protein expression did not exhibit corresponding variations in protein levels. A likely explanation in the case of HDL is that, for some proteins, only a limited amount of the protein can be incorporated into the HDL lipid-protein complex, the remainder presumably being degraded.

**Table 3.** Correlations between HDL proteins and clinical traits measured within HMDP

| **Trait1** | **Trait2** | **Bicor Value** | **P-value** |
| --- | --- | --- | --- |
| Apoc3 | HOMA-IR (pre-bleed) | 0.50 | 0.0001 |
| Apoc1 | Liver collagen | 0.49 | 0.0011 |
| Ttr | LiverResidual | 0.49 | 6.67E-06 |
| Ihh | Insulin (pre-bleed) | 0.47 | 0.0005 |
| Plg | lesion area | 0.46 | 0.0002 |
| Ighm | lesion area | 0.46 | 0.0003 |
| Gm5938 | lesion area | 0.45 | 0.0004 |
| Apoc3 | Insulin (pre-bleed) | 0.44 | 0.0015 |
| Ihh | HOMA-IR (pre-bleed) | 0.44 | 0.0012 |
| Apoc2 | Kidney (left) | 0.43 | 0.0001 |
| Gc | unesterified cholesterol | 0.43 | 0.0002 |
| Obp1a | lesion area | 0.42 | 0.0012 |
| Alb | LiverResidual | 0.38 | 0.0006 |
| Mup4 | RBC | 0.38 | 0.0007 |
| Apoc3 | triglycerides (pre-bleed) | 0.38 | 0.0009 |
| Mup4 | HGB | 0.36 | 0.0014 |
| Gc | VLDL+LDL | 0.36 | 0.0016 |
| Gc | total cholesterol | 0.35 | 0.0017 |
| Apoc2 | PLT | 0.34 | 0.0025 |
| Pon3 | free fluid | -0.41 | 0.0002 |
| Acta2 | HOMA-IR | -0.40 | 0.0003 |
| Apoh | MONO | -0.37 | 0.0011 |
| Ihh | free fluid | -0.36 | 0.0014 |
| Fetub | MCV | -0.35 | 0.0020 |
| Scgb2b7 | adiposity | -0.35 | 0.0024 |
| Acta2 | Insulin | -0.34 | 0.0024 |
| Apoa1 | average fat mass (liver) | -0.34 | 0.0025 |
| Psap | GRAN % | -0.34 | 0.0026 |
| Apoa5 | WBC Time | 0.34 | 0.0027 |
| Acta2 | Insulin | -0.34 | 0.0024 |
| Psap | GRAN % | -0.34 | 0.0026 |
| Antxr1 | GRAN % | -0.34 | 0.0027 |
| Gc | triglycerides (liver) | -0.34 | 0.0040 |
| Ighm | free fatty acids (pre-bleed) | -0.34 | 0.0035 |

We have attempted to identify distal (*trans*-acting) factors affecting HDL protein levels (Supplemental Table 2). In contrast to local eQTL, where only SNPs within 1Mb of the gene are tested for association, distal QTL analyses involve genome wide SNP tests for association, requiring a much higher threshold for significance. A number of the likely significant distal eQTL occur with several Mb of the gene and, thus, probably result from either long range (>1 Mb) linkage disequilibrium or chromosome looping interactions (for example *Apoh, Apom, B2m, H2-Q10,* and *Pp1c*) (Supplemental Table 2). The pQTL analysis also identified some highly significant distal (*trans*-acting) interactions, most notably for *Apoa2* (p=4.6 x 10^−14^). The *Apoa2* locus is about 5 Mb from the structural gene and the significant association is probably the result of linkage disequilibrium or chromosome looping.

We asked if the local pQTL could be used to identify causal interactions between HDL proteins. For this, we selected the genes with significant pQTL and asked whether the peak local pQTL SNP was associated with any other HDL proteins, suggesting that the regulation of the former perturbs levels of the latter. For example, *Apoe3* (on chr. 9) protein levels were controlled by a local pQTL with peak SNP 46673334 (chr. 9) and the same SNP was significantly associated with the levels of *Podx1* (chr. 6, p=4×10^−3^), *Fetub* (chr. 16, p=5.5×10^−3^), *Apoc2* (chr. 7, p=5.9×10^−3^), and a number of other proteins. Likewise, a local pQTL SNP for *Apoa2* was associated with *Apoc3* and *Itgb3* levels and an *Hq-Q10* local pQTL SNP was associated with *Apob* and *Apoe* levels (Supplemental Table 4).

**Table 4.** The peak QTLs for sterol efflux capacity of HDL. The peak phenotypic QTL locus with p < 10^-6 are listed along with SNP identifiers and their locations

| SNP | CHR | BP | P | Beta |
| --- | --- | --- | --- | --- |
| **Baseline Efflux** |  |  |  |  |
| rs30557586 | 3 | 55764730 | 4.13E-07 | 8.19E-01 |
| rs31424282 | 3 | 57945327 | 1.74E-06 | -8.38E-01 |
| rs29379333 | 10 | 69242755 | 6.21E-06 | 8.73E-01 |
| rs6333057 | 12 | 58206555 | 5.43E-07 | 8.76E-01 |
| rs50224465 | 12 | 53464782 | 8.55E-07 | 1.07E+00 |
| rs31810918 | 15 | 37462413 | 2.74E-08 | 9.80E-01 |
| rs3718535 | 16 | 9212865 | 1.65E-06 | 7.18E-01 |
| rs6284288 | 18 | 4398494 | 8.11E-06 | 6.97E-01 |
| rs13483883 | 20 | 93699907 | 5.59E-06 | 8.40E-01 |
| **Stimulated Efflux** |  |  |  |  |
| rs30557586 | 3 | 55764730 | 6.61E-06 | 6.68E-01 |
| **ABCA1 Dependent Efflux** |  |  |  |  |
| rs31551612 | 1 | 171208377 | 2.75E-06 | -5.86E-01 |
| rs45738488 | 6 | 67533600 | 4.43E-06 | -6.30E-01 |
| rs31630237 | 8 | 118112906 | 2.04E-07 | -6.28E-01 |
| rs26919597 | 11 | 68048005 | 6.78E-06 | -6.30E-01 |
| rs46923442 | 13 | 5984274 | 1.40E-07 | -7.33E-01 |
| rs47306105 | 13 | 6601243 | 2.27E-06 | -7.10E-01 |
| rs51267071 | 13 | 6240479 | 4.24E-06 | -6.80E-01 |
| rs46153864 | 13 | 6257029 | 4.24E-06 | -6.80E-01 |
| rs50517602 | 13 | 6322879 | 4.24E-06 | -6.80E-01 |
| rs49326176 | 13 | 6416532 | 4.24E-06 | -6.80E-01 |
| rs47351631 | 13 | 6520213 | 4.24E-06 | -6.80E-01 |
| rs46431216 | 13 | 6087220 | 4.48E-06 | -6.82E-01 |
| rs6373590 | 13 | 6229579 | 4.48E-06 | -6.82E-01 |
| rs37828224 | 13 | 8785486 | 9.10E-06 | -6.93E-01 |
| rs36839806 | 19 | 57424289 | 8.34E-07 | -7.01E-01 |

### Clustering of HDL metrics based on quantitation across strains

To understand interactions of HDL proteins with each other and with other metrics of HDL (ABCA1 specific sterol efflux, baseline-diffusion sterol efflux, HDL particle size, and HDL cholesterol), we correlated proteins that are present in more than 80% of the strains. We then applied hierarchical clustering to a matrix that contains all measured phenotypes. The clustering of the proteome and functional metrics revealed expected patterns (Figure 3 for yeast normalized and Supplemental Figure 3 for total normalized data). The complex correlation matrices represented a high number of strongly correlated variables suggesting an organized interplay among HDL proteins and between the physiological and functional metrics of HDL (Figure. 4 and Supplemental Figure 4 for yeast and total normalized data respectively). Of the 8100 total correlations, we have focused on 2216 correlations that are |r|>0.5 with a Bonferroni-Holm adjusted P<0.05, n=2216. The correlation and *P* values are presented in Supplemental Table 2.

**Figure 3.**
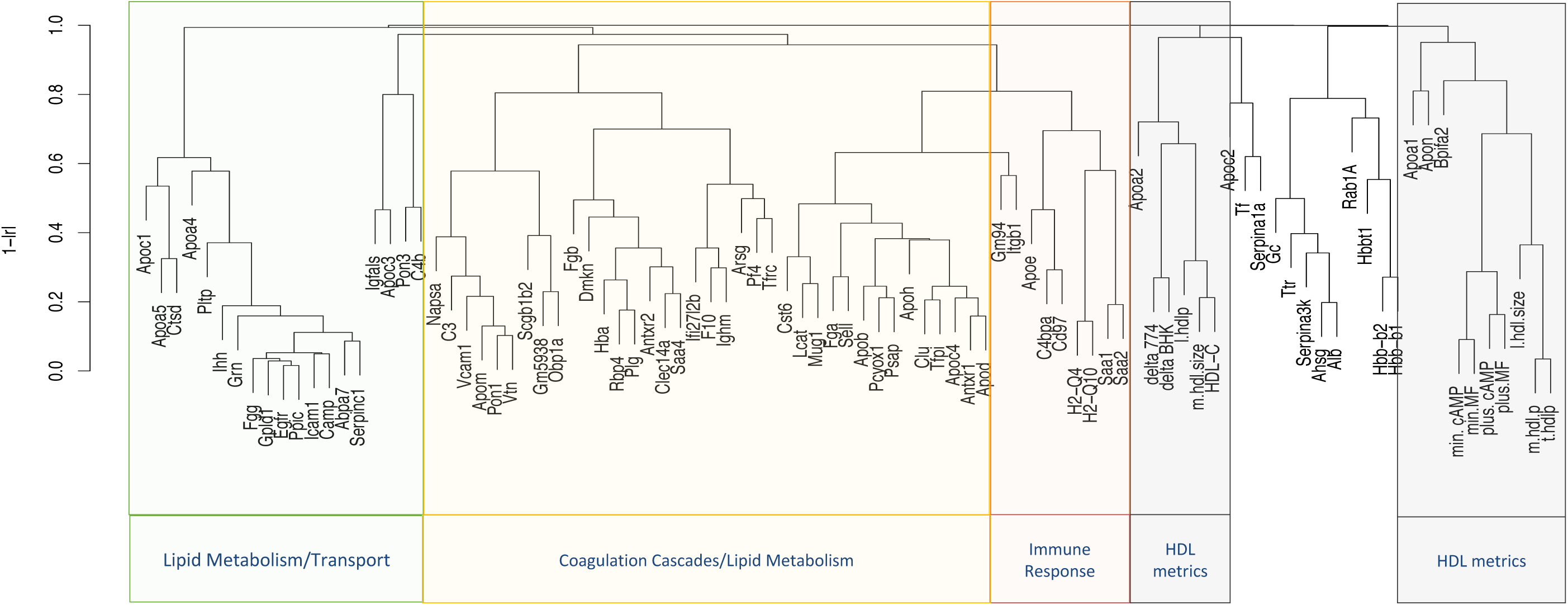
Hierarchical clustering of the HDL metrics: Proteome, sterol efflux, particle concentration and size. min.MF, plus.MF, delta.BHK are unstimulated, ABCA1 upregulated and ABCA1 specific sterol efflux from BHK cells respectively. min.cAMP, plus.cAMP, delta.J774 are unstimulated, ABCA1 upregulated and ABCA1 specific sterol efflux from J774 cells respectively. m.hdl.size and l.hdl.size are medium and large HDL sizes respectively. The correlation structure was determined using pearson correlation. The protein functional groups were curated from DAVID, KEGG, Panther, and Uniprot databases.

**Figure 4.**
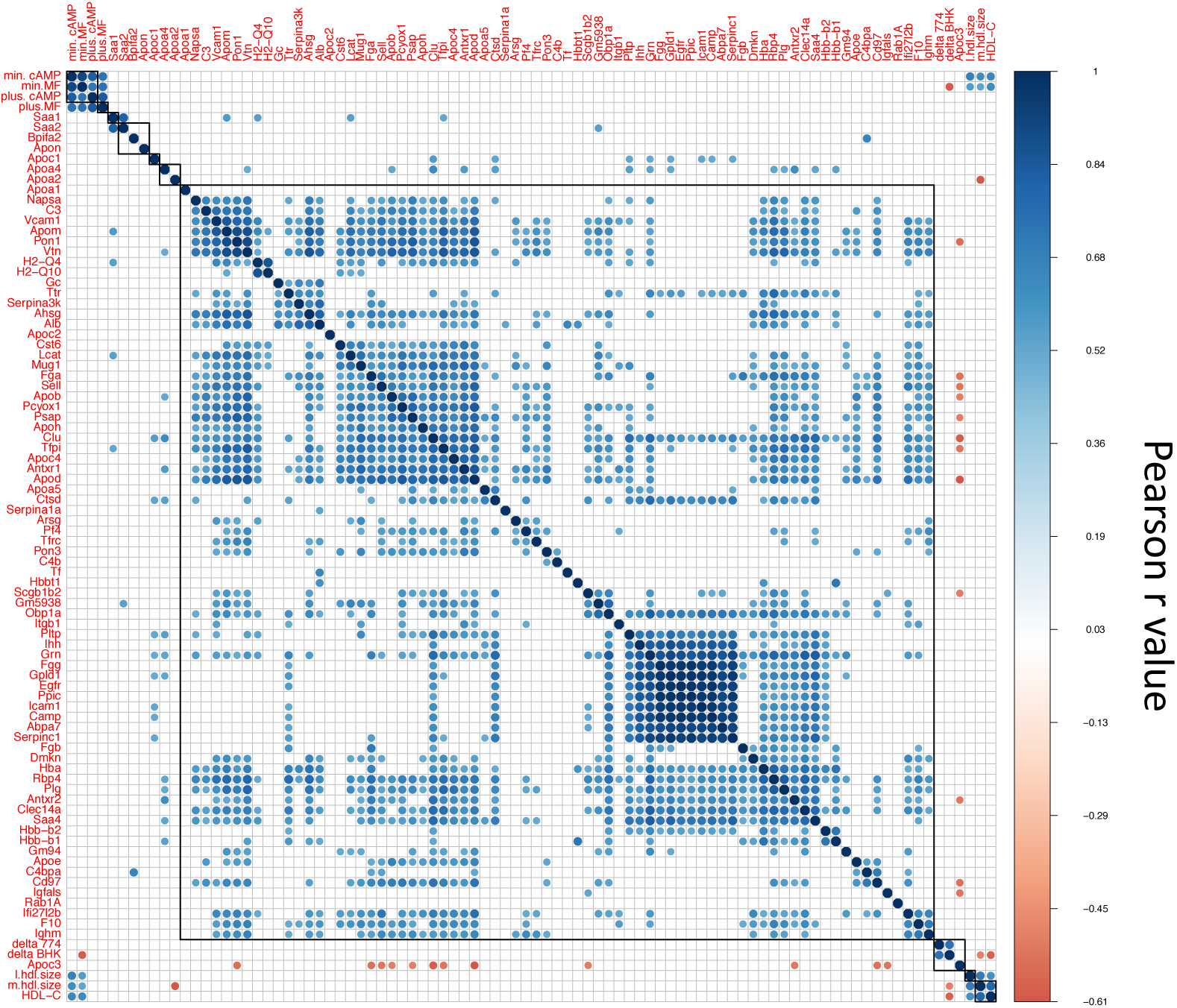
The relationship between HDL metrics is represented by a correlation matrix. Total of 8,100 correlations were observed. Among which 2,216 Pearson correlation with Bonferroni-Holm correction having |r| values >0.5 (positive in bleu, negative in red) that are *P*<0.05 are presented.

All sterol efflux metrics displayed up to 4-fold differences across strains (Supplementary Figure 7). The non-proteome phenotypes clustered together; for example, unstimulated sterol efflux from two different cell types, min.cAMP and min.MF and stimulated sterol efflux (r= 0.909, *P*=10^−15^), plus. cAMP and plus.MF (r=0.83, *P*=10^−15^) correlated strongly and were part of the same cluster that is modestly associated with APOA1 (Figure 3 and Supplementary Table 2). The ABCA1 mediated sterol efflux, delta BHK and delta J774, correlated strongly (r=0.73, *P*=10^−15^) and formed a cluster along with HDL-C and medium size HDL where the latter two metrics correlated strongly (r=0.78, *P*=1.82e-12) (Figure 3 and Supplementary Table 5).

HDL-C was negatively associated with ABCA1 mediated sterol efflux from both BHK and J774 cells (r=-0.58, *P*=0.00036 and −0.49, *P=*0.03 respectively) and was positively associated with the diffusional (unstimulated) efflux from both cell types (r=0.62 P=6.14e-06 for BHK and r=0.64 P=2.04e-05 for J774).

Mice have 85% of their HDL cholesterol distributed in the 7.6-9.8nm range (medium size) in a monodispersed peak (Pamir *et al*, 2016), and therefore medium size HDL represents the majority of HDL cholesterol. This latter cluster is associated with APOA2 levels (Figure 3). Further, proteins that participate in immune responses, including serum amyloids SAA1, SAA2, histocompatibility complexes H2-Q4 and H2-Q10 associated strongly and formed a distinct cluster with other immune response proteins CD97, C4BPA and APOE. All of the hemoglobin proteins, HBBA, HBB1, HBBT1 formed a distinct cluster. Lipid metabolism proteins, APOC1, APOA5, APOA4, PLTP, and GPLD1 clustered together. Most interestingly, APOC3 correlated negatively with 39 proteins on HDL. Among these were proteins with roles in immune response such as APOE, PON1, SAA1, SAA4 (Supplementary Table 5). insulin like growth factor binding protein, IGFALS clustered strongly but negatively with APOC3 (r=-0.53, *P*=0.002). Recent studies suggest a role for APOC3 in beta cell insulin resistance (Åvall *et al*, 2015) and according to proteome interactome by Harmonizome (a tool curated from 100 public databases (Rouillard *et al*, 2016)) APOC3 is one of the 73 proteins found to interact with IGFALS.

We observed significant correlations between HDL-C levels and GM94, ITGB1, APOC1, APOC2 (positive) and APOA2, SERPINA1A, ALB, and TFRC (negative) regardless of PSM normalization method used (Supplemental Table 5). All the New Zealand strains studied (NZB BIN/J, NZW LAC/J, KKHI/J) were in the top quintile of HDL-C distribution and bottom quintile of APOA2 distribution. The QTL analysis indicated the same genomic regions for regulation of APOA2 and HDL-C levels (Supplemental Figure 8). APOA2 seems to be the main genetic regulator of plasma HDL-C levels in mice.

Among the HDL proteins, APOD, with high homology to carrier proteins such as lipocalins and with strong innate immune response roles such as antioxidative (Ganfornina et al., 2008) and neuroprotective effects (Do Carmo et al., 2008) had the most, significant correlations with other HDL proteins involved in immune response. Twenty of these correlations exhibited |r|>0.7 suggesting that it is highly interactive apolipoprotein that acts as a carrier for other proteins on HDL.

Another way to visualize the relationships among the proteins is to present these as a correlation network (Figure 5 and Supplemental Figure 5). The network consists of multiple layers of spatial organization, a core, an outer layer and the periphery. The “core” proteins, including AHSG, NAPSA, PLG, SAA4, HBA, APOD, APOJ, APOM, APOH, FGA, and TFPI are surrounded by common HDL associated apolipoproteins such as APOA4, APOA5, APOC1, APOC3, APOC4 in addition proteins that have been described to be associated with HDL in most proteomic studies (Davidson HDL proteome watch http://homepages.uc.edu/∼davidswm/HDLproteome.html) such as H2-Q4, H2-Q10, SELL, ANTRX1/2, PF4 RBP4, SERPINs, GPLD1, FGA and FGB. More peripheral are known associated proteins such as PLTP, PON3, LCAT, APOE, SAA1, SAA2. HDL core proteins formed a tight network suggesting that they are co-regulated. Inflammation response and complement activation proteins such as SAA1, SAA2, H2-Q4, H2-Q10, C3, C4b, C4BPA were distal to the core co-regulatory network suggesting that they are mostly regulated by external factors such as an inflammatory stimulus. The two major structural proteins, APOA1 and APOA2 along with APOC3 were on the outer layers and periphery and are negatively correlated with other proteins suggesting that their presence requires the displacement of other proteins.

**Figure 5.**
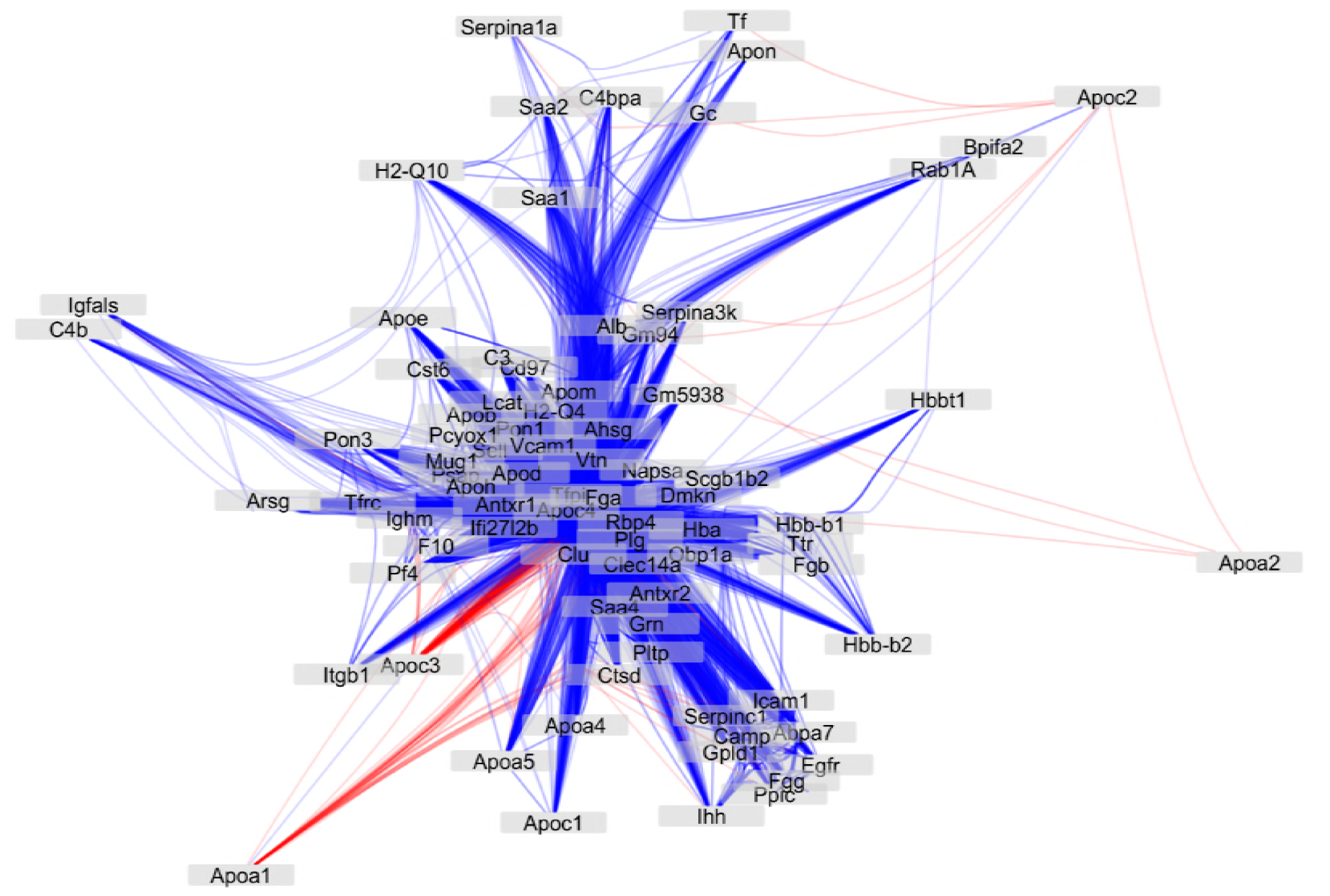
Cytoscape visualization of all the protein-protein interactions presented in Figure 1. Self-loops were removed and edges were bundled for clarity. Node locations are assigned using an edge-weighted spring-embedded layout algorithm using the negative log of the Benjamini-Hochberg corrected p-value, and edge transparency is directly proportional to the same value. Red: negative correlation, Blue: positive correlation. Shorter distances indicate stronger correlations.

### Genetic Regulation of Sterol Efflux

The sterol efflux capacity was measured in two different cell lines (J774 and BHK) in basal (no ABCA1) and stimulated (with ABCA1) conditions, with ABCA1 specific sterol efflux calculated by subtracting basal from stimulated. The QTL analysis for the efflux traits revealed shared global association profiles that are condition specific and in agreement between cell types (Supplemental Figure 6 A and B). The significant QTLs that are shared by both cell types are presented in Table 4. The lack of strong QTLs that associate with sterol efflux capacity of HDL suggest that either sterol efflux is under limited genetic control or its control is multigenic with small effect sizes.

### Lipid and clinical trait interactions

The HMDP strains have been examined in separate studies (Lusis et al., 2016) either on chow (Bennett et al., 2010), high fat/high sucrose (Parks et al., 2013; 2015) or high cholesterol diets (Bennett et al., 2015)2. We compared our data with results from these studies, reasoning that while power to detect correlation will be diminished given that the data were collected in separate animals and at different times with incomplete overlap of strains, traits significantly impacted by genetics should still retain some correlation structure. The most significant correlations between HDL proteins and clinical traits are summarized in Table 3 and detailed in Supplemental Table 6.

A total of 22 HDL proteins significantly and positively correlated with lesion area in the hypercholesterolemia study (Bennett et al., 2015) among which Plasminogen (PLG), Immunoglobin chain C region (IGHM), Platelet Factor 4 (PF4) were strongest (r=0.469, P=0.00026; r=0.461, P=0.00034, r=0.37, P=0.0049). We have recently shown that plasminogen is an effective sterol acceptor through ABCA1 and this could be the mechanism by which plasminogen contributes to atherosclerosis(Pamir et al., 2017).

Conforming to the recently identified role of APOC3 in insulin resistance, HOMA-IR plasma insulin levels, body weight and adiposity correlated positively with HDL associated APOC3 levels (r=0.50, P=0.00017; r=0.44 P=0.0015; r=0.23, P=0.048; r=0.26, P=0.021, respectively).

## Discussion

We report the analysis of the HDL proteome across a set of 93 inbred strains of mice exhibiting common genetic variation. The genetic variation across this mouse panel resembles that in human populations, based on the number of common SNPs (about 4 million). Previous studies in mice have revealed genetic variations in the levels of HDL-cholesterol, HDL apoproteins, and HDL composition (Lusis et al., 1983). Our results are consistent with a high level of heritability of HDL proteins, including the identification of a number of novel pQTLs. One significant conclusion that has emerged is that certain proteins cluster together in response to genetic perturbations, presumably reflecting physical or regulatory interactions. These clusters then help define the heterogeneity of HDL particles, although our analyses do not address lipid heterogeneity. Another important conclusion is evidence of a relationship between protein composition and HDL function. Finally, we have identified potential links between HDL proteins and various clinical or molecular traits studied previously in the HMDP strains. We discuss each of these points in turn below.

Normalization of shotgun proteomic data is a continuous struggle in the field (Välikangas et al., 2018). HDL is a rather uncomplicated mixture containing only ∼100 proteins. However, its proteome is driven by the 10 most abundant proteins, with 65% being made up of APOA1 and 15% of APOA2 (Toth et al., 2013). The normalization strategy should conserve the compositional bias of the HDL. The normalization that directly adjusts scale such as Total Count (TC) and Upper Quartile (UQ) fails to accommodate compositional bias. The normalization strategies that adjust scales using landmarks in the distribution Median, (Med), Differentially expressed (DESeq), Trimmed mean (TMM) are promising approaches for HDL proteome however, detailed analyses need to be performed for their validation to be used on smaller libraries of stochastic count data from mass spectrometry. The quartile (Q) and reads per kilobase per million mapped (RPKM) (equivalent of normalizing spectral counts to protein length) have adverse effects on intra-sample variance and on distribution bias (Dillies et al., 2013). The TC and UQ normalizations favor the most abundant proteins and are unfriendly for mixtures with a distribution bias. That said, TC normalization is often the preferred method for shotgun HDL proteomics as it controls for differences in instrument response, digestion efficiency, and amounts of loaded protein digest but fails at conserving the distribution bias as it tends to accommodate the changes in the abundant proteins(Vaisar et al., 2007). That is partly why the HDL protein quantification by shotgun proteomics is not optimal and the correlation with immunobased assays are moderate (Hoofnagle et al., 2012). Therefore, we opted to include a second normalization approach by spiking yeast carboxy peptidase at levels ∼8 fold lower than APOA1 and to correspond to the median/mean abundance of the typical HDL proteome (Carvalho et al., 2008; Liu et al., 2004). The QTLs identified using both proteomic information are mostly overlapping with TC normalization resulting in ∼30% more significant QTLs. The relationships among inbred lines of mice was inferred from the high-density SNP map where strains cluster according to their genealogy (Cervino et al., 2005). We employed the same approach: The 155 proteins that were present in at least 20% of the strains loosely predicted the relatedness of the strains according to their genealogy for inbred strains and according to the breeding scheme for recombinant strains. Almost half of these proteins (81) were present in greater than 80% of the strains and only 34 were shared by all the strains. The strain dependent distribution of HDL proteome across 93 strains validates our previous studies with only 5 strains (Pamir et al., 2016). However, the comparison of the clustering patterns between microarray data or the SNPs did not reach full agreement as genetic variation explains only a fraction of the variation and a very small part of the genome is involved in regulating HDL (data not shown). The 93 strains are represented by N=1-5 with a distribution of ∼4, 9, 75, 9, and 1 % for N=1,2,3,4, and 5 respectively. Even though, these N are not optimal to calculate – intra and –inter strain variation, the broad sense heritability calculations captured 65 proteins that have greater than 10% heritability. Among which APOA2 has a score of 0.62 which is consistent with its strong association with HDL-C loci – a highly heritable trait.

The high-level heritability of the proteins is demonstrated by >20K pQTL associated SNPs that map to > 66 loci. To understand whether a pQTL results from structural or regulatory variation, we have incorporated gene expression information. A positive finding in such analysis suggests that the genotype dependent differential gene expression is the basis of most of the association. (Farber et al., 2011). In our studies, we used adipose and liver tissue global gene expression profiles to map distinct loci in liver (5) and adipose tissue (20) and that are associated with the gene expression levels for the SNPs associated with pQTLs. Adipose tissue exhibited only inflammatory gene (*Saa1, Saa2, Tfrc, Vtn*) associated SNPs that were almost exclusively *trans-* acting. Liver tissue had *cis* and *trans* eQTLs associated with multiple genes including *Apoc4, Apoh, Fgb, Tfrc, Saa2*. The complex regulation at the protein and gene expression level dictates the protein composition of HDL and its hereditary preservation.

Correlation networks such as weighted gene co-expression network analysis is a systems biology method for describing the correlation patterns among genes. The weighted correlation network analysis (WGCNA), revealed that core HDL proteins, composed of most common apoproteins are highly correlated and co-regulated. The co-regulated gene network is consistent with HDL’s role in innate immunity and lipid metabolism as it reveals a tight network of co-regulation among the proteins with primary roles in immunity and lipid metabolism. The histocompatibility protein isoforms such as H2Q4 and H2-Q10 that have been shown to be associated with mouse HDL in multiple studies (S. M. Gordon et al., 2010; 2015; Pamir et al., 2016) are part of the core co-regulated proteins. In mice, histocompatibility proteins play a role in innate immunity by antigen presenting via major histocompatibility complex class 1 (MHC-1).H2-Q10 is the only murine MHC-1 protein found in the serum in appreciable concentrations (Lew et al., 1986). While these innate immune proteins with roles in antigen presentation are part of core co-regulation network, acute phase proteins such SAA1 and SAA2 are not, as they are primarily regulated by an inflammatory stimulus.

While up to 50% of HDL cholesterol level can be heritable, less is known about heritability of its sterol efflux function or its proteome (Bentley et al., 2014; Goode et al., 2007). The sterol efflux capacity of HDL seems to be regulated by a multigenic architecture with small effect size. The loci captured using 93 strains of mice have moderate P values and small effect sizes. In a human cohort of 846 individuals, Villard et al tested 7 preselected SNPs with known effects in HDL metabolism such as ABCA1, CETP, APOA1 and APOA2. Seven SNPs tested accounted together for approximately 6% of total plasma efflux capacity supporting our findings of moderate strength QTLs.

The classic linear view of HDL genesis from discoidal, lipid-poor nascent particles to spherical, cholesterol- and phospholipid-rich particles packed with a combination of over 100 different proteins has been recently challenged by the finding that HDL is secreted directly from hepatocytes in 4 distinct sizes, with little interchange between them, and representing all of the plasma HDL subparticle pools (Mendivil et al., 2016). Although our analyses do not incorporate the lipidome of HDL which can contribute to the orchestration of the composition of HDL sub populations, and our samples weren’t stored with a cryoprotectant (HDL structure, function and proteome can be affected and proteome under represented (Holzer *et al*, 2017) a highly inter correlated proteome reveals the complexity of HDL particle composition. We captured a remarkable 2216 correlations among the proteins that survived multiple comparison correction and that explains at least 25% of the variation (R2>0.25). The hierarchical clustering of the correlated proteins regrouped the proteins according to their biological functions emphasizing the coordinated co-regulation. However, while applying the stringent statistical approach certain biological relationships were missed. For example, APOC3 significantly and exclusively negatively correlated with 36 other HDL proteins (Supplementary Table 2). In humans, }increased circulating APOC3 levels are associated with cardiovascular disorders, inflammation, and insulin resistance (Chan et al., 2008; Petersen et al., 2010). On the other hand, humans with an APOC3 mutation benefit from a favorable lipoprotein profile, increased insulin sensitivity, lower incidence of hypertension, and protection against cardiovascular diseases (Atzmon et al., 2006; Jørgensen et al., 2014; Pollin et al., 2008). The negative correlation of APOC3 with 36 other proteins, its association with plasma insulin levels, and HOMA-IR levels conforms to its newly appreciated role as a brake on the metabolic system. Efforts to identify the proteomic, lipidomic, and functional fingerprints of HDL subspecies are of critical importance and may open paths to novel pharmacological targets.

Clinical and epidemiological studies show a robust, inverse association between HDL cholesterol (HDL-C) levels and CHD risk (D. J. Gordon and Rifkind, 1989; Wilson et al., 1988). However, pharmacological interventions aimed at raising HDL cholesterol levels in humans showed no cardiovascular benefits (AIM-HIGH Investigators et al., 2011; Barter et al., 2007; Landray et al., 2014; Schwartz et al., 2012). Since the collapse of the HDL-C hypothesis for atherosclerosis, a new generation of HDL metrics are under investigation to be used in clinic (Fazio and Pamir, 2016). For example, greater HDL cholesterol efflux capacity (CEC), independent of levels of HDL-C and APOA1 (the major structural protein of HDL), is associated with a lower prevalence of atherosclerotic vascular disease (Khera et al., 2011; Li et al., 2013; Rohatgi et al., 2014). Most changes in HDL function are likely to be a reflection of changes in the HDL proteome (Green et al., 2008; Vaisar et al., 2007). Thus, identification of the protein signature responsible for loss of sterol efflux capacity could provide biomarkers of clinical validity to assess CHD risk. The interplay between HDL sterol efflux function, particle concentration, size and HDL proteome is still poorly understood. HDL-C levels correlated strongly with all the efflux measures. While we captured strong associations between expected metrics such as diffusional or ABCA1 specific efflux from two different cell types, no single HDL protein explained the majority of the variation in sterol efflux, suggesting that it is a polygenic process. That said, APOA2 explained about 10% of ABCA1 dependent sterol efflux from both cell types and the *Apoa2* locus was strongly associated with ABCA1 specific sterol efflux (data not shown). In mice, APOA2 seems to impact the sterol efflux function at the protein and gene level. It is important to note that *Apoa2* locus aligns with HDL-C QTL.

In summary, a systems biology approach reveals the highly complex and inter-correlated nature of HDL protein composition, heritable contributions to both HDL levels and composition and its association with disease. We show that HDL proteins preserve hereditary patterns that are likely to harbor ancestral/lineage information. It is likely that inheritance controls the production of HDL particles of a certain protein and lipid composition that have different functions. At present, we lack a model for the assembly of HDL protein and lipid cargo. Our results provide the ground work to support future studies aimed at characterization of the genetic architecture regulating HDL function and comprehensive composition in humans.

## Methods

### Mice

All studies were approved by the Animal Care and Use Committee of the University of California, Los Angeles. Mice were housed (1-3/cage) in a pathogen-free barrier facility (22 °C) with a 12 h light/dark cycle with free access to food and water. All the strains were a fed low-fat diet (Wayne Rodent BLOX 8604; Harlan Teklad Laboratory). 60-80 day old mice fasted for 16 h at 7pm and sacrificed at 9am the following morning. Mice were bled from the retro-orbital sinus into tubes containing EDTA (final concentration 1 mM), after isofluorane inhalation. Plasma was collected and stored at −80 °C until analysis.

### Plasma HDL-C measurements

Plasma cholesterol levels (Invitrogen) were determined biochemically following the manufacturer’s guidelines.

### Cholesterol efflux assays

Macrophage cholesterol efflux capacity was assessed with J774 macrophages labeled with [^3^H]cholesterol and stimulated with a cAMP analogue, as described by Rothblat, Rader, and colleagues (la Llera-Moya et al., 2010). Efflux via the ABCA1 pathways were measured with BHK cells expressing mifepristone-inducible human ABCA1 that were radiolabeled with [^3^H]cholesterol (Shao et al., 2005). Efflux of [^3^H]cholesterol was measured after a 4 h incubation in medium with APOB depleted serum HDL (2.8% v/v). ABCA1-specific cholesterol efflux capacity was calculated as the percentage of total [^3^H]cholesterol (medium plus cell) released into the medium of BHK cells stimulated with mifepristone after the value obtained with cells stimulated with medium alone was subtracted.

### HDL isolation

Serum HDL was prepared by adding calcium (2 mM final concentration) to plasma and using PEG (polyethylene glycol, 8kDa, Sigma) to precipitate lipoproteins containing APOB (VLDL, IDL, LDL). After centrifugation at 10,000 g for 30 min at 4 °C, serum HDL was harvested from the supernatant. HDL was isolated from serum or EDTA-anticoagulated plasma, using sequential ultracentrifugation (d=1.063-1.21 mg/mL) (Vaisar et al., 2007). HDL was stored on ice in the dark and used within 1 week of preparation. For each isolation batch, control samples from the same pooled mouse plasma were included and further processed by tryptic digest and MS to control for experimental variability.

### Liquid chromatography-electrospray ionization tandem mass spectrometric (LC-ESI-MS/MS) analysis

HDL (10 μg protein) isolated by ultracentrifugation and 0.5 ug of yeast carboxypeptidase were solubilized with 0.1% RapiGest (Waters) in 200 mM ammonium bicarbonate, reduced with dithiothreitol, alkylated with iodoacetamide, and digested with trypsin (1:20, w/w HDL protein; Promega) for 3 h at 37 °C. After a second aliquot of trypsin (1:20, w/w HDL protein) was added, samples were incubated overnight at 37 °C. After RapiGest was removed by acid hydrolysis, samples were dried and stored at −20 °C until analysis. Prior to analysis, samples were reconstituted in 5% acetonitrile, 0.1% formic acid (Pamir et al., 2016). Tryptic digests of mouse HDL (1 μg protein) were injected onto a C18 trap column (Paradigm Platinum Peptide Nanotrap, 0.15 x 50 mm; Michrom Bioresources Inc.), desalted (50 μL/min) for 5 min with 1% acetonitrile/0.1% formic acid, eluted onto an analytical reverse-phase column (0.15 x 150 mm, Magic C18AQ, 5 μm, 200 A; Michrom Bioresources Inc.), and separated on a Paradigm M4B HPLC (Michrom Bioresources Inc.) at a flow rate of 1 μL/min over 180 min, using a linear gradient of 5% to 35% buffer B (90% acetonitrile, 0.1% formic acid) in buffer A (5% acetonitrile, 0.1% formic acid). Electrospray ionization (ESI) was performed using a Captive Spray source (Michrom BioResources, Inc, Auburn, CA) at 10 mL/min flow rate and 1.4 kV setting. HDL digests were introduced into the gas phase by ESI, positive ion mass spectra were acquired with a orbitrap mass spectrometer (Fusion, Thermo Electron Corp.) using data-dependent acquisition (one MS survey scan followed by MS/MS scans of the 8 most abundant ions in the survey scan) with a 350-1400 *m/z* scan. An exclusion window of 30 s was used after 2 acquisitions of the same precursor ion (Pamir et al., 2016; Vaisar et al., 2007).

### Protein identification

MS/MS spectra were matched against performed using the Comet search engine (version 2015.01 rev. 1) against a mouse UniProt database appended with yeast caboxypeptidase Y protein sequence (52,639 total sequences). The following Comet search parameters were applied: peptide mass tolerance of +/−20.0 ppm allowing for C13 isotope offsets, full tryptic digest allowing up to 2 missed cleavages, oxidized methionine variable modification, and carbamidomethyl cysteine static modification. The search results were subsequently processed though the Trans-Proteomic Pipeline (version 4.8.0) using the PeptideProphet and ProteinProphet tools using an adjusted probability of ≥0.90 for peptides and ≥0.95 for proteins. Each charge state of a peptide was considered a unique identification (Nesvizhskii et al., 2003). We used the gene and protein names in the Entrez databases (National Center for Biotechnology Information; based on the nomenclature guidelines of the Human Gene Nomenclature Committee (http://www.gene.ucl.ca.uk/nomenlature) for human (Wain et al., 2002) and Mouse Genome Informatics (http://www.infromatics.jax.org.nomen/) guidelines (Davisson, 1994) to identify HDL proteins and to eliminate the redundant identifications of isoforms and protein fragments frequently found in databases used in proteomic analysis (Rappsilber and Mann, 2002). This approach also permits cross-referencing of proteins from different species.

### Protein quantification

Proteins were quantified using Peptide Spectra Matches (PSM) —the total number of MS/MS spectra detected for a protein (Vaisar et al., 2007). Proteins considered for analysis had to be detected in ≥30 analyses (%10 of the total samples) with ≥2 unique peptides. Because only a few peptides are typically measured for a given protein, these peptides might not be sufficient to define all isoforms of the protein that are present in the sample therefore, when MS/MS spectra could not differentiate between protein isoforms, the isoform with the most unique peptides was used for further analysis.

PSMs for each protein, normalized to either spiked yeast carboxypeptidase or to total PSMs for peptides from each sample, were used to calculate a normalized PSM to compare the relative protein composition of mouse strains’ HDLs (Vaisar et al., 2007). Supplemental Table 1 provides the total calculated PSMs for each protein, the individual peptides that identified each protein, the total number peptide spectra matches, and relative quantification as normalized to yeast carboxypeptidase Y total PSMs or total PSMs of one sample.

### HDL particle size

HDL particle size was quantified by calibrated ion mobility analysis (Hutchins et al., 2014). Briefly, HDL isolated by ultracentrifugation from EDTA plasma is introduced into the gas-phase ions by ESI. Because electrophoretic mobility depends chiefly on size, ion mobility analysis data are expressed in terms of particle diameter (nm), which corresponds to the calculated diameter of a singly charged, spherical particle with the same electrophoretic mobility (Hutchins et al., 2014).

### Association analyses

GWAS for protein levels and gene expression was performed using correction for population structure as described (Hui et al., 2015; Orozco et al., 2012). Loci were defined as *cis* if the peak SNP mapped within 1 Mb of gene position and *trans* if it mapped outside (*cis* significance threshold, p < 1.4 e-3; *trans* threshold, p < 6.13e-6).

### Heritability

Broad sense heretibiliaty scores were calculated for each protein using R package (sommer) using the formula H2= genetic variance / (genetic variance + residual variance)

### Statistical analyses

Data are means ± SEMs. Linear correlation among the HDL metrics of the 93 strains were assessed with pearson correlations and the association of the proteins were assessed by spearman correlations both were followed with Bonferroni-Holm post-hoc correction for multiple comparisons. Data were analyzed with Prism and R software.

### Data and software availability

The MS/MS datasets produced in this study are available in the PRIDE consortium (ProteomXchange submission ref: 1-20180406-10713) and in the UCLA based public database established to harbor HMDP related data (https://systems.genetics.ucla.edu/data).

## Acknowledgements

**Source of Funding**

This work was supported by awards from the National Institutes of Health and American Heart Association. HL112625, HL108897, P30 DK17047, P01 HL092969, T32HL007828, HL076491, HL30568 (AJL); HL122677 and HL28481, SDG18440015(T.Q.de A.V.); SDG16940064 (NP); HL121214 (CT); HL112625, HL108897, P30 DK17047, P01 HL092969, T32HL007828, HL076491. P01 HL092969, P30 DK017047 (JWH)

## Disclosures

Jay Heinecke is named as a co-inventor on patents from the US Patent Office on the use of HDL markers to predict the risk of cardiovascular disease. Dr. Heinecke has served as a consultant for Kowa, Merck, Amgen, Bristol Meyer Squibb, GSK, and Pacific Biomarkers.

## Figure Legends

**Supplemental Figure 1:**
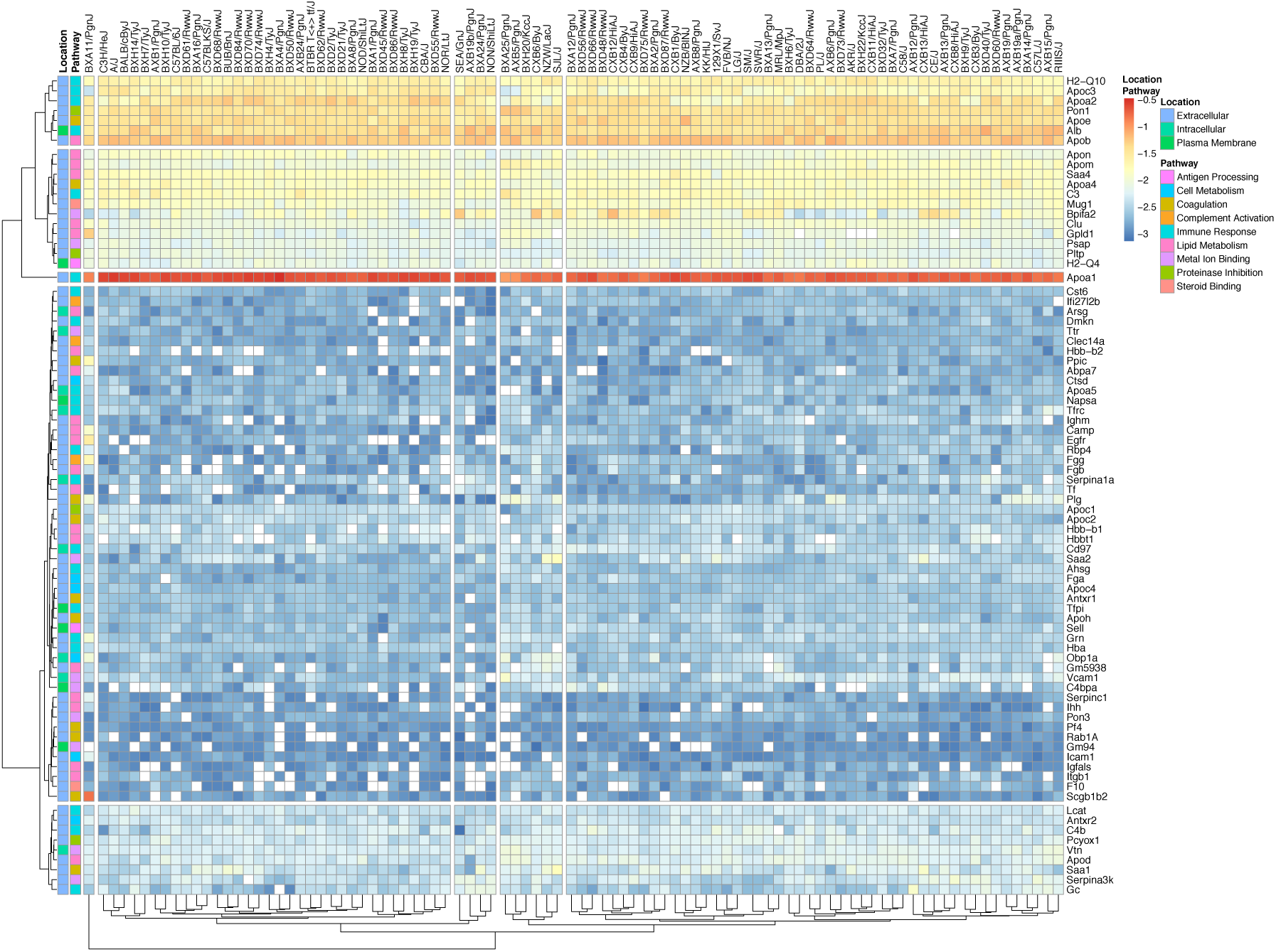
The heatmap visualization of the HDL protein abundances (calculated as normalized to total PSMs) across 93 strains. The proteins, their biological functions and cellular locations are represented. Logarithmic transformation of the total PSM normalized data has been performed to accommodate the abundance distribution form high (red) to very low (dark blue). White squares represent not available values. Both the proteins and the strains were clustered using Euclidean distances.

**Supplementary Figure 2.**
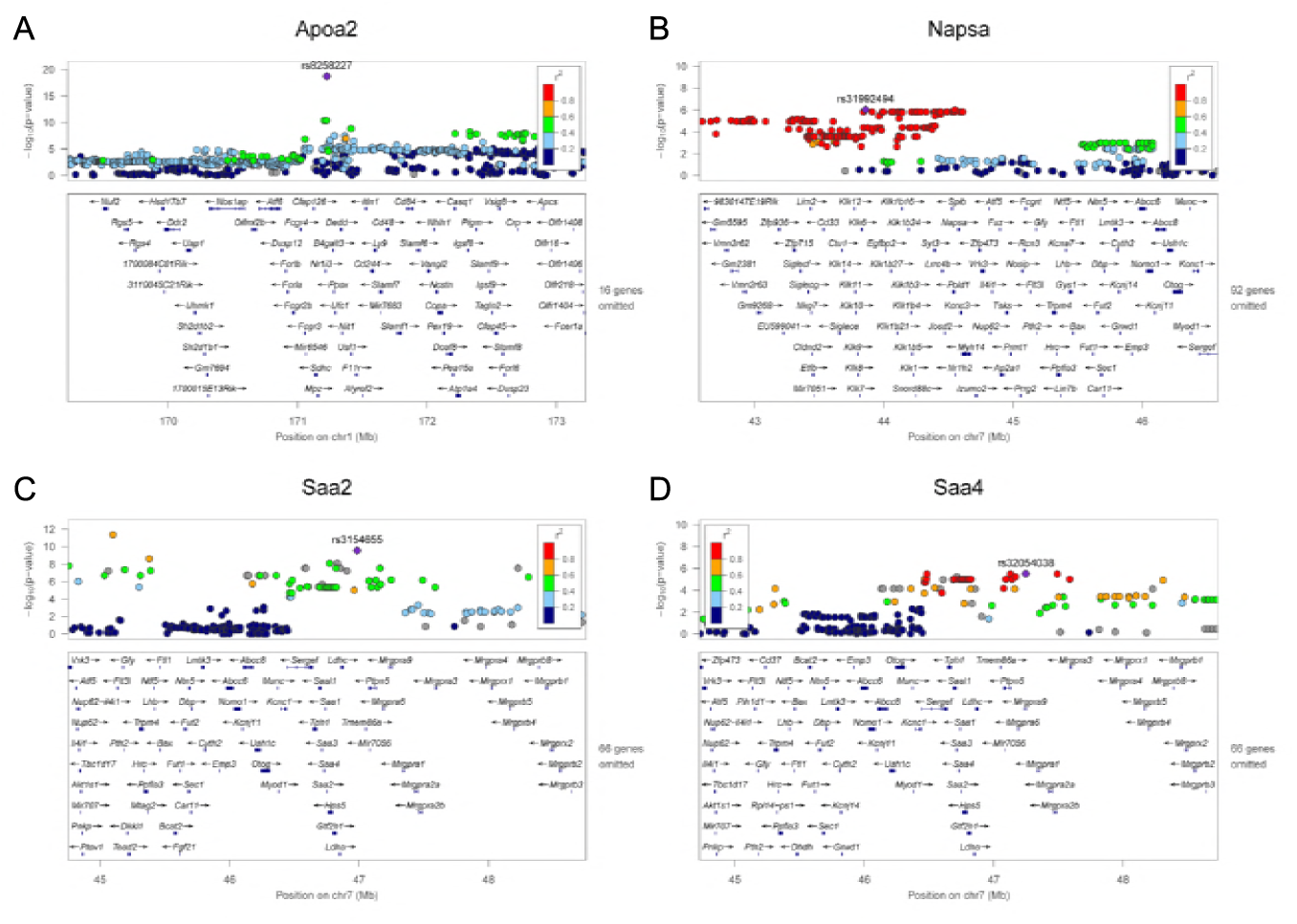
A representative QTLs for HDL proteins normalized to total PSMs. Loci associated with A) APOA2, B), NAPSA, C) SAA2, D) SAA4.

**Supplementary Figure 3.**
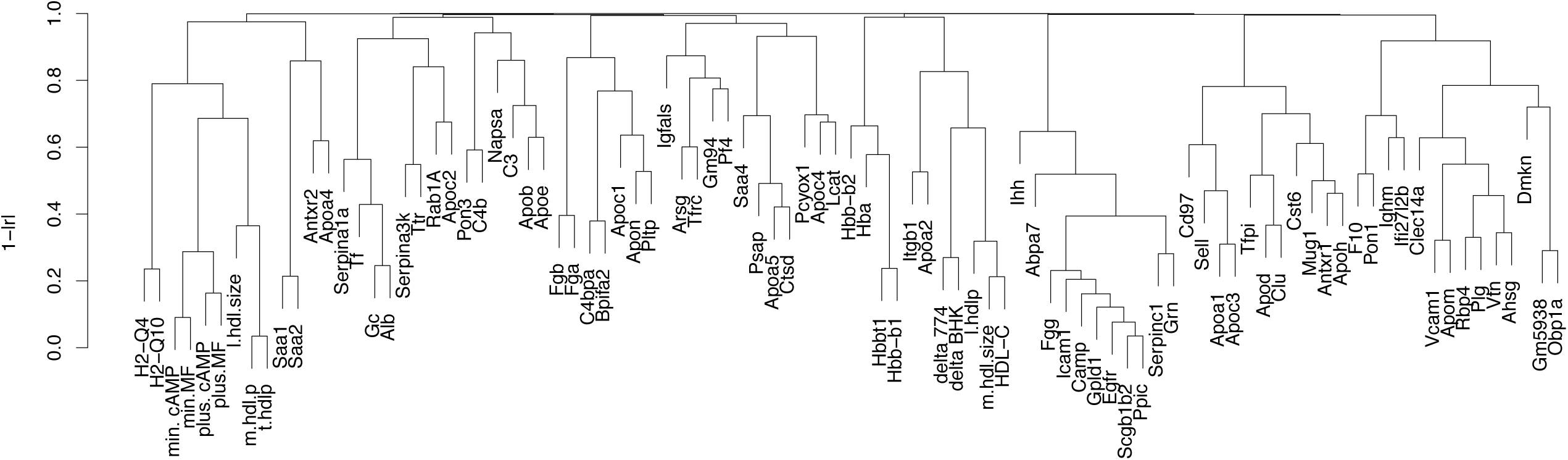
Hierarchical clustering of the HDL metrics: Proteome (normalized to total PSMs), sterol efflux, particle concentration and size. min.MF, plus.MF, delta.BHK are unstimulated, ABCA1 upregulated and ABCA1 specific sterol efflux from BHK cells respectively. min.cAMP, plus.cAMP, delta.J774 are unstimulated, ABCA1 upregulated and ABCA1 specific sterol efflux from J774 cells respectively. m.hdlp and t.hdlp are medium size and total HDL particle concentration. m.hdl.size and l.hdl.size are medium and large HDL sizes respectively. The correlation structure was determined using pearson correlation. The protein functional groups were curated from DAVID, KEGG, Panther, and Uniprot databases.

**Supplementary Figure 4.**
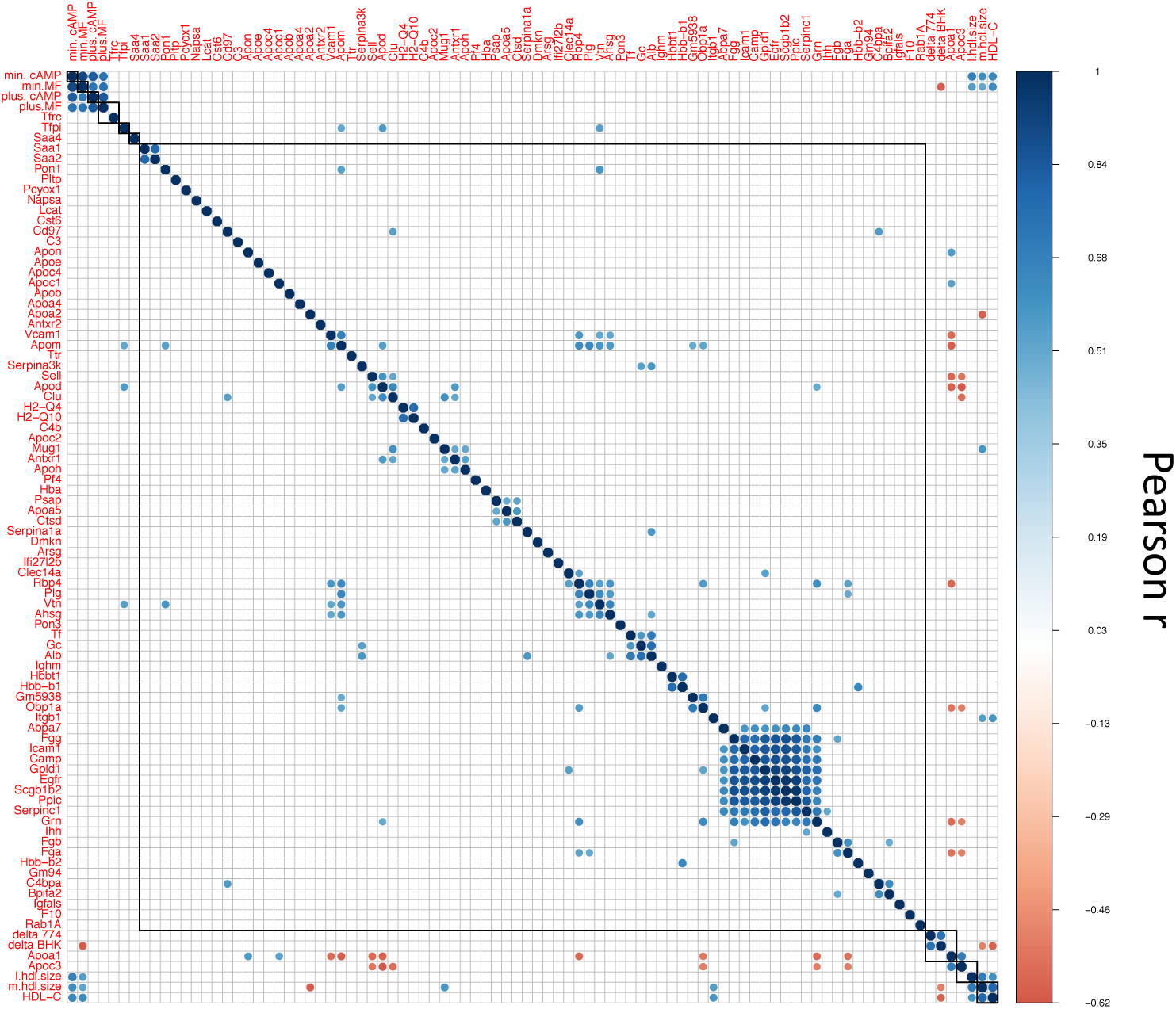
The relationship between HDL metrics is represented by a correlation matrix. Total of 8,100 correlations were observed. Among which 380 Pearson correlation with Bonferroni-Holm correction having |r| values >0.5 (positive in bleu, negative in red) that are *P*<0.05 are presented.

**Supplemental Figure 5.**
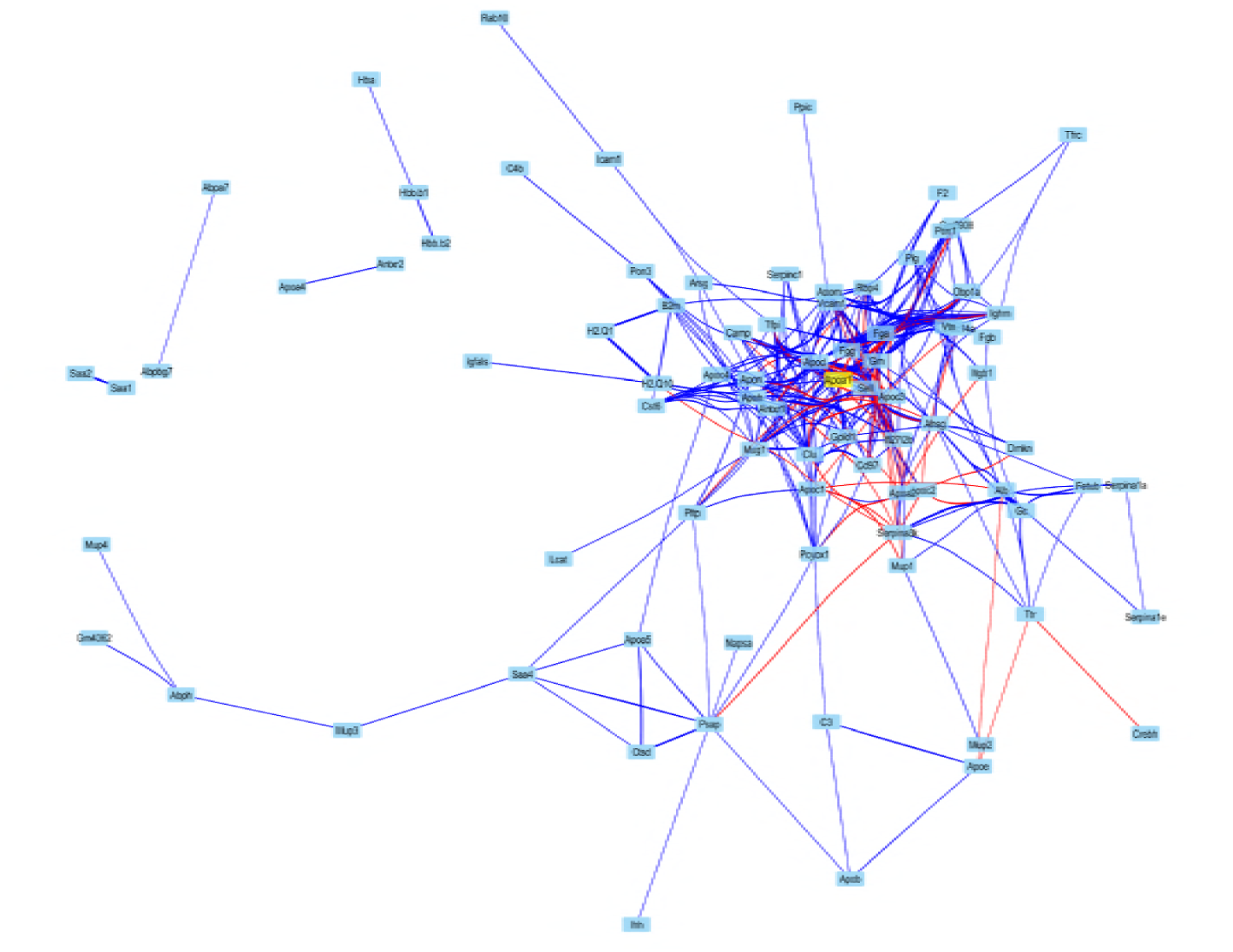
Cytoscape visualization of all the protein-protein interactions presented in Supplemental Figure 2. Self-loops were removed and edges were bundled for clarity. Node locations are assigned using an edge-weighted spring-embedded layout algorithm using the negative log of the Benjamini-Hochberg corrected p-value, and edge transparency is directly proportional to the same value. Red: negative correlation, Blue: positive correlation. Shorter distances indicate stronger correlations.

**Supplemental Figure 6.**
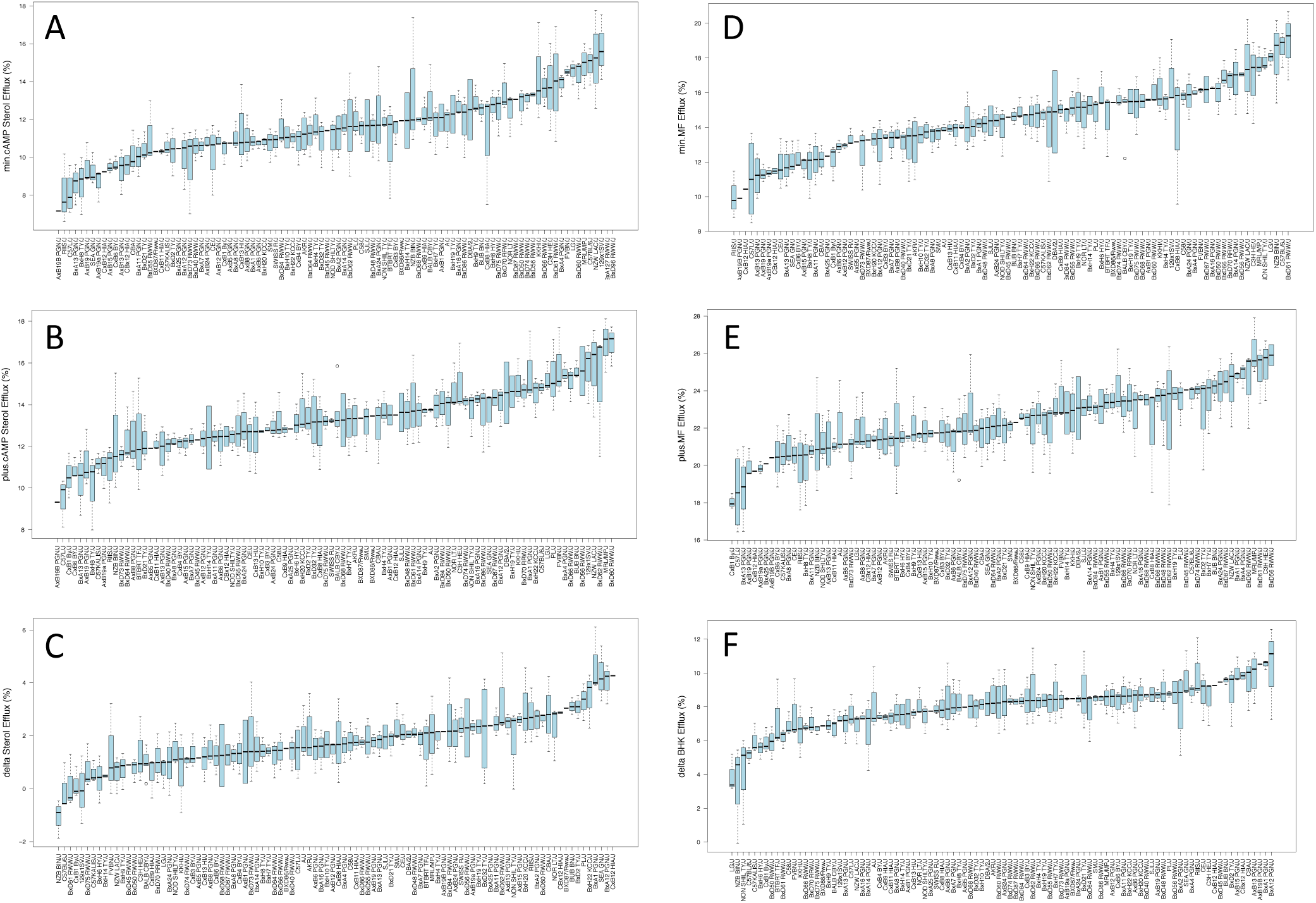
Sterol Efflux distribution across strains. min.cAMP, plus.cAMP, delta.J774 are unstimulated, ABCA1 upregulated and ABCA1 specific sterol efflux from J774 cells respectively (-C). min.MF, plus.MF, delta.BHK are unstimulated, ABCA1 upregulated and ABCA1 specific sterol efflux from BHK cells respectively (D-F). All the measures are in duplicate from PEG depleted plasma collected from 1-5 mice for each strain.

**Supplemental Figure 7.**
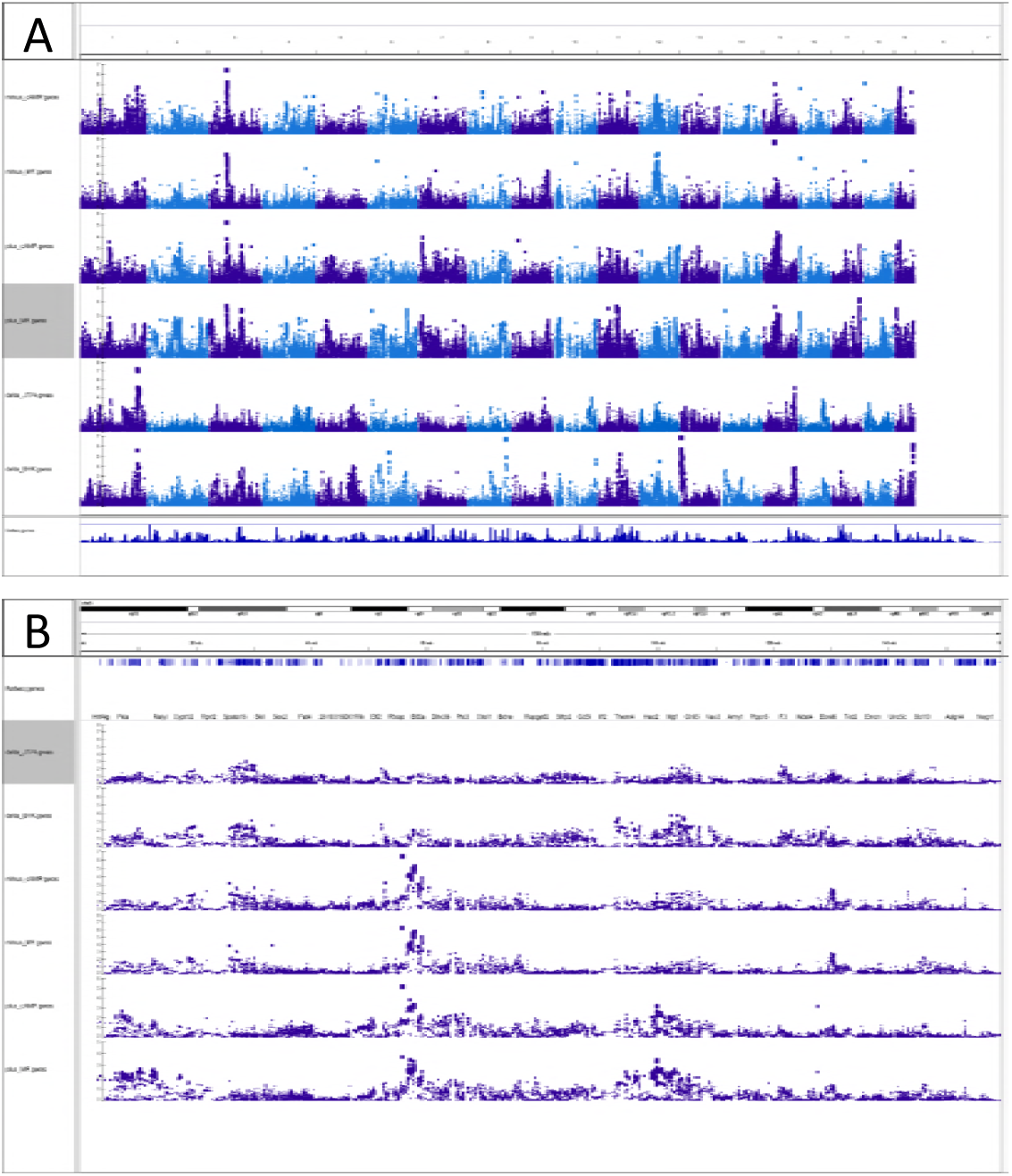
**The manhattan plots showing A) Global association profiles across strains for baseline (min. BHK**, min.cAMP) stimulated (plus.BHK, plus.cAMP) and ABCA1 dependent (delta.J774 delta.BHK) sterol efflux; B) The sterol efflux QTLs located on chromosome 3.

**Supplemental Figure 8.**
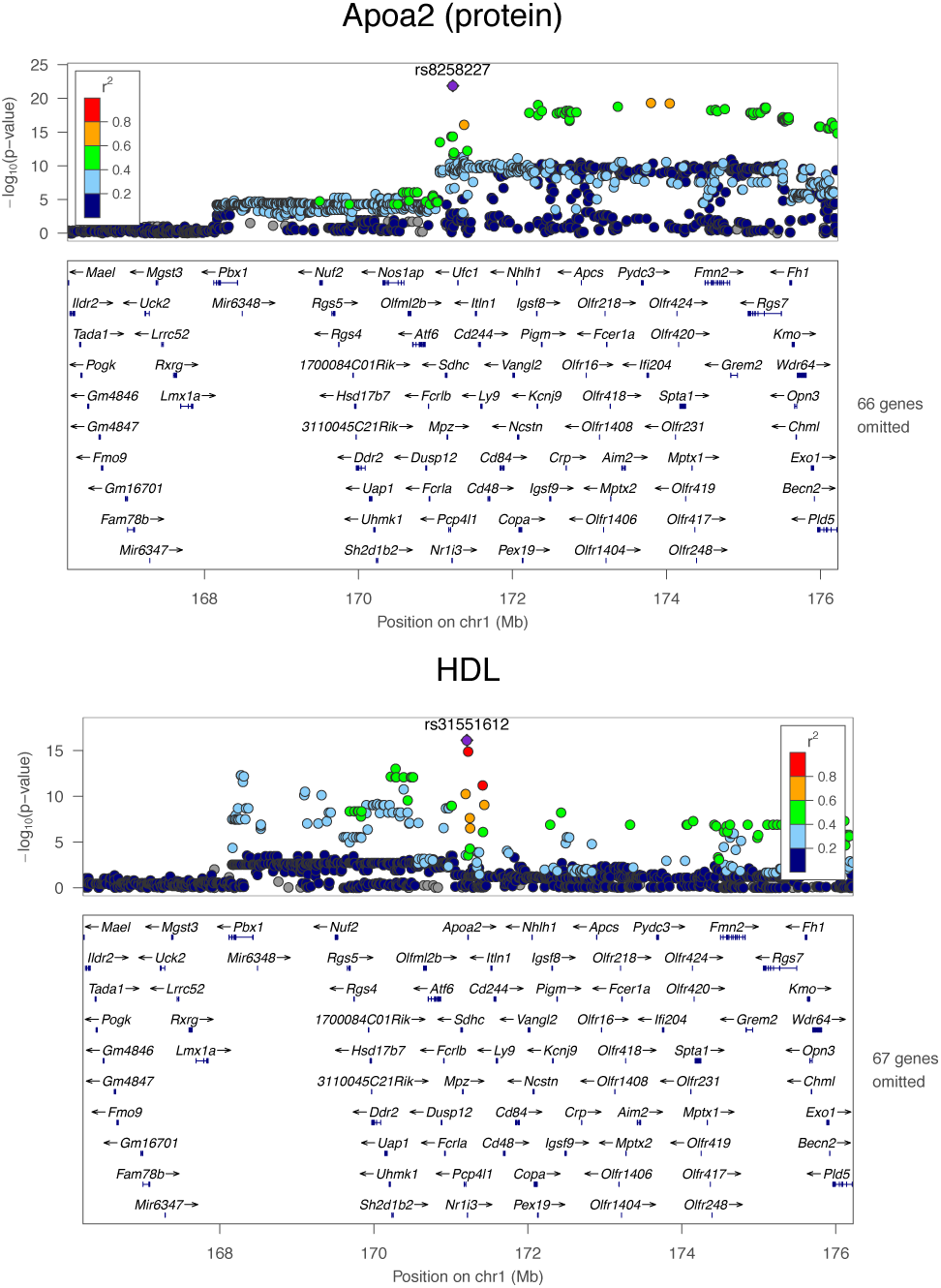
Manhattan plot showing alignment between APOA2 QTL and HDL-C QTL. Overlap of the loci regulating APOA2 and HDL-C.

**Supplementary Table 1.** Summary of HDL proteins per strain.

**Supplemental Table 2.** Distal hepatic eQTL and distal HDL pQTL.

**Supplemental Table 3.** Local adipose eQTL and local HDL pQTL.

**Supplemental Table 4.** Potential causal interaction based on local pQTL analyses.

**Supplementary Table 5.** The correlations for the HDL metrics.

**Supplemental Table 6.** Association of HDL Proteins with Clinical Traits in HMDP.

